# Active mesh and neural network pipeline for cell aggregate segmentation

**DOI:** 10.1101/2023.02.17.528925

**Authors:** Matthew B. Smith, Hugh Sparks, Jorge Almagro, Agathe Chaigne, Axel Behrens, Chris Dunsby, Guillaume Salbreux

## Abstract

Segmenting cells within cellular aggregates in 3D is a growing challenge in cell biology, due to improvements in capacity and accuracy of microscopy techniques. Here we describe a pipeline to segment images of cell aggregates in 3D. The pipeline combines neural network segmentations with active meshes. We apply our segmentation method to cultured mouse mammary duct organoids imaged over 24 hours with oblique plane microscopy, a high-throughput light-sheet fluorescence microscopy technique. We show that our method can also be applied to images of mouse embryonic stem cells imaged with a spinning disc microscope. We segment individual cells based on nuclei and cell membrane fluorescent markers, and track cells over time. We describe metrics to quantify the quality of the automated segmentation. Our segmentation pipeline involves a Fiji plugin which implement active meshes deformation and allows a user to create training data, automatically obtain segmentation meshes from original image data or neural network prediction, and manually curate segmentation data to identify and correct mistakes. Our active meshes-based approach facilitates segmentation postprocessing, correction, and integration with neural network prediction.

**Statement of significance:** In vitro culture of organ-like structures derived from stem cells, so-called organoids, allows to image tissue morphogenetic processes with high temporal and spatial resolution. Three-dimensional segmentation of cell shape in timelapse movies of these developing organoids is however a significant challenge. In this work, we propose an image analysis pipeline for cell aggregates that combines deep learning with active contour segmentations. This combination offers a flexible and efficient way to segment three-dimensional cell images, which we illustrate with by segmenting datasets of growing mammary gland organoids and mouse embryonic stem cells.

## I. INTRODUCTION

We describe here a full pipeline for segmenting microscopy images of cells in 3D, using active meshes and artificial neural networks. This includes a plugin for Fiji, Deforming Mesh 3D (DM3D), which provides an assisted way to segment cells in 3D over time. We apply our pipeline to segmentation of dynamic, relatively small cell aggregates (*∼* 10’s of cells).

The field of segmenting and tracking cells and nuclei in 3D microscopy images has experienced numerous recent developments [1]. Semi-automated or assisted tools such as ilastik [1] or Labkit [3] can be used to segment images using pixel classification. Leveraging neural networks, techniques such as StarDist [4] allow the users to generate segmentations automatically, in the case of StarDist by localizing nuclei using star-convex polygons. In these tools, segmentations can be obtained by either using a pretrained model, or creating training data manually and training a new model, or by augmenting an existing model through generating new training data and further training. Other tools that use neural networks are Cellpose [5], which creates a topological map where gradient flow tracking [6] is used to find the contour of the cell, and Embed-Seg [7], an embedding-based instance segmentation method. These techniques are appropriate for detecting and segmenting cells as binary blobs. Another technique to segment cells involves creating a mesh representation and evolving active contours to best fit the image [8–10]. Integrating tracking with detection can improve segmentation efficiency, as tracking algorithms or networks can be used to predict cells in successive frames and improve the seeding of new cells for segmentation [11–16].

Our technique uses a workflow common to other neural-network based methods: the user can manually segment a subset of data, then use a neural network to automatically create more segmentations for the remaining data. Our method however incorporates the use of active meshes in this workflow for initial manual segmentation, for automatically segmenting the neural network generated images, and for manual correction. This brings an important advantage, as editing meshes in 3D is an intuitive and convenient way to perform 3D segmentation, notably compared to using 2D pixel based segmentation tools. Active meshes are handled and deformed using a custom-made Fiji plugin, Deforming Mesh 3D (DM3D). This plugin is based on an implementation of an active mesh deformation method and handles several segmentation meshes in the same image frame.

In our pipeline (Fig. 1), manually obtained 3D meshes are used to create labels that are learned by a neural network with a 3D Unet architecture [17]. One of the labels the neural network learns to create is the distance transform, a label which associates to each voxel a value corresponding to its distance to the edge of the object it is associated with. The distance transform or watershed transform [18] have been used previously in combination with deep learning neural networks for object detection and separating overlapping objects [18, 19].

**FIG. 1:**
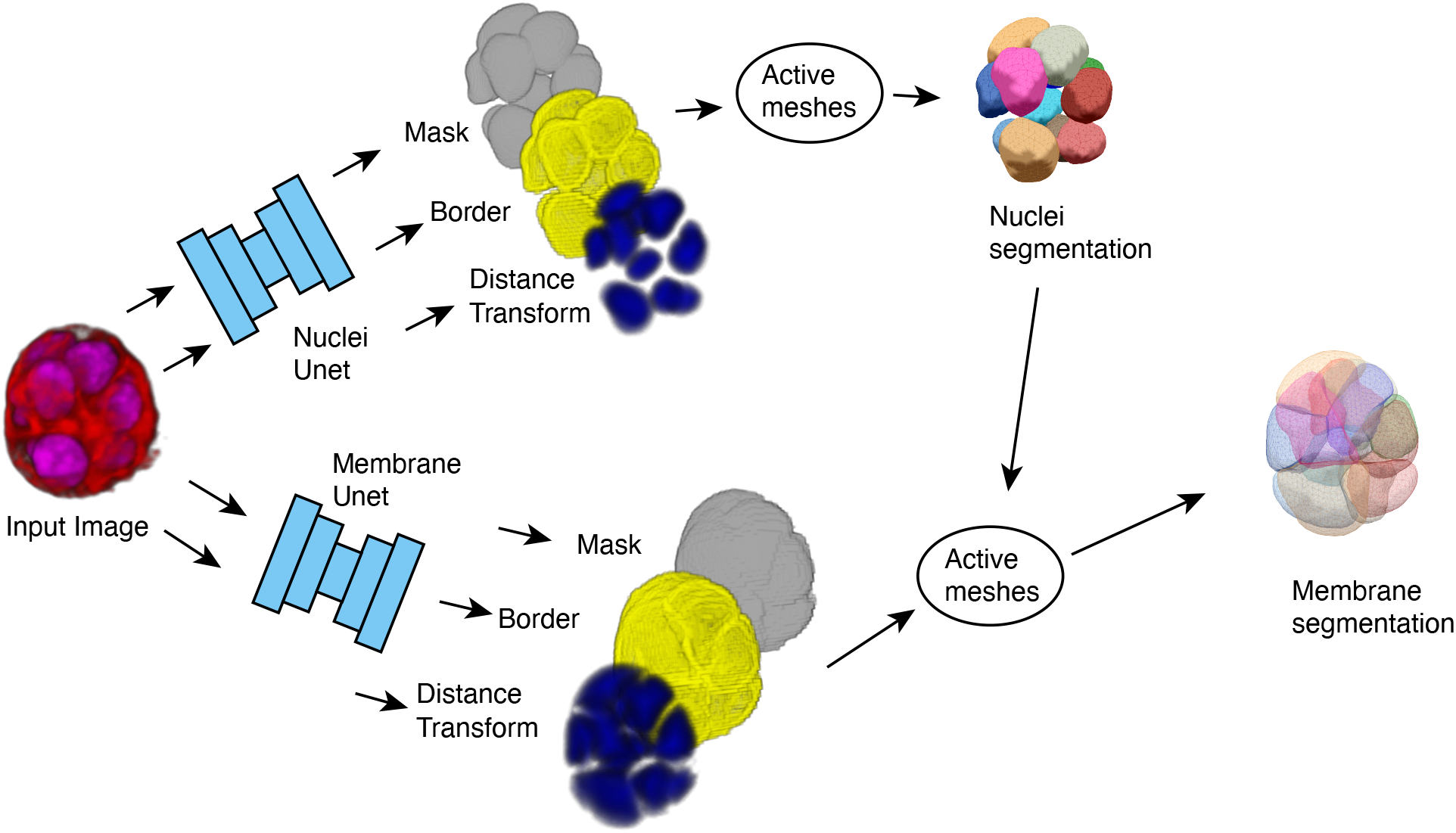
Overview of segmentation pipeline, from an original 2-channel 3D fluorescent microscopy image to a set of meshes that represent the cell nuclei and the cell membranes.

The trained neural network processes a 3D timelapse movie and predicts a modified distance transform for each voxel within each frame. The distance transform is modified in the sense that it takes non-zero values only within the surface which it measures the distance from. This distance transform is used to locate 3D regions that represent individual cells or their nuclei. A triangulated mesh is initialized within each of these regions. An active mesh method is then used to deform the mesh to the outer surface of nuclei or cell membranes.

To demonstrate the effectiveness of our technique we segmented and tracked 6 mammary gland organoids for 24 hours at 11 minutes imaging intervals (Fig. 2). Organoids have nuclei labelled with the dye SiR-DNA and membrane labelled with tdTomato (see Material and Methods). Image data was obtained using multichannel dual-view oblique plane microscopy [20], and we selected organoids that appeared to have good signal to noise at the beginning of the imaging period. We refer to this dataset as Movies 1-6.

**FIG. 2:**
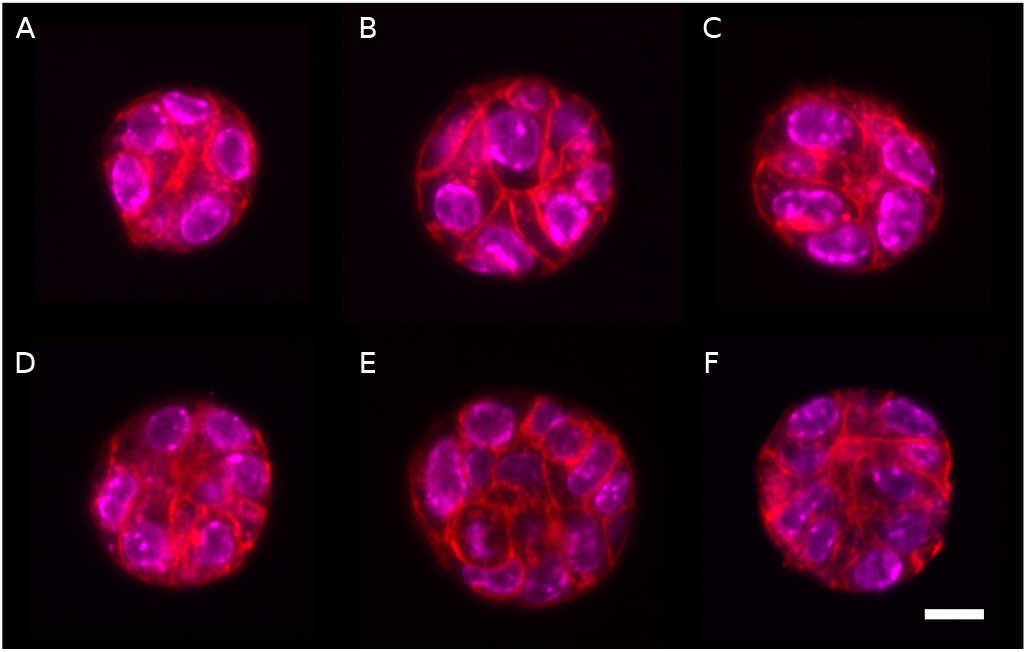
x-y cross sections through the equator of 6 different organoids after 8 hours of imaging. Scale bar is 10 µm. Red label: membrane dye, magenta: DNA label. Organoids in A-F are later referred to as Movies 1-6.

To segment this dataset, we first generated original training data by manually creating segmentations of a subset of the data. We then processed the whole dataset with a trained neural network to obtain initialisation for segmentation meshes, which are deformed using the DM3D plugin. We then refined the generated segmentations by manual inspection and tracking cells with DM3D, to segment the complete timelapse movies.

To evaluate the quality of the neural network segmentations, we prepared ground truth data set from manual segmentations and compared that to segmentations from the fully automated pipeline. We show an overview of the segmentation results, and a measure of their quality, by comparing results from the pipeline to manual segmentations.

To also verify that our pipeline can be applied to different types of cells and microscopy images, in section IV we also quantify segmentation results of mouse embryonic stem cells imaged with a spinning disc microscope.

## II. METHODS

### A. Manual segmentation of original image data

Here we describe the mesh-based segmentation technique we use to manually segment cell nuclei and cell membranes from original image data (Fig. 3). To generate manual segmentation using DM3D we initialise a coarse version of the nucleus or the cell to segment in 3D. This is performed by manually positioning spheres within the nucleus or the cell, trying to capture their shape. A mesh approximating the shape of the resulting collection of spheres is created, using a raycast technique to fill the spheres [8]. This initial mesh is subsequently deformed to conform to the nucleus shape, by minimizing an effective energy with two contributions: an intrinsic force which depends on the mesh shape as described in Appendix B 1, and a force arising from an “image energy” that depends on the mesh and on the voxel values. We use different effective energies for manually segmenting nuclei and cell membranes from original image data, as described below.

**FIG. 3:**
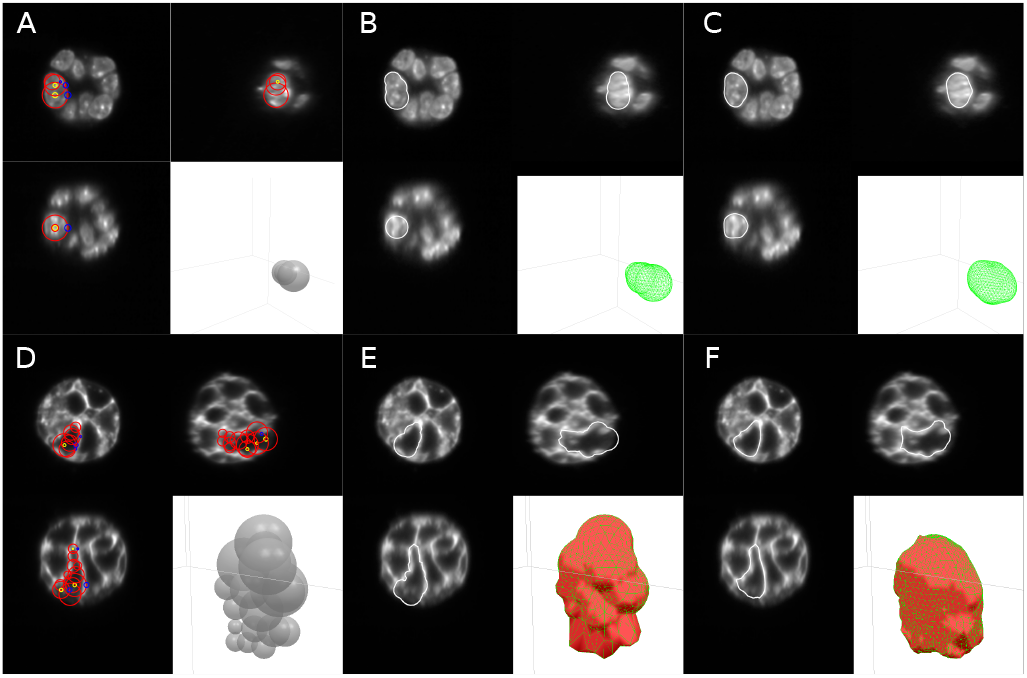
Manual initialisation of segmentation meshes that are then deformed using the active mesh method to the cell nucleus (A-C) or to the cell membrane (D-F). A,D) Orthogonal cross section views and a 3D view during mesh initialization. Red circles: boundaries of the spheres used for mesh initialization. The yellow and blue circles are handles that can be manipulated by the user to adjust the position and radius of the spheres. B, E) Same orthogonal views with the initialized mesh. F,G) Mesh after deformation to the nucleus or cell membrane image intensity.

#### 1. Segmentation of cell nuclei from original image data

To deform meshes to outer surfaces of nuclei, we use a “perpendicular gradient energy”. Labelled nuclei are essentially 3D-filled continuous regions of high intensity. Therefore, we use an an energy that is based on the gradient of the nuclear channel [21]. We denote *I*(**x**) the image intensity at a voxel position **x**. We associate a unit normal vector **n** to a node on the mesh by averaging and normalising the unit normal vectors to triangles connected to the node. The energy associated to a node on the mesh and evaluated at position **x** is then defined as:

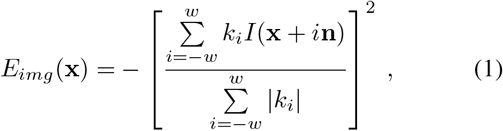

with 𝒩is a normalisation factor, we choose *w* = 5, and the coefficients *k*_*i*_ are obtained from the derivative of a Gaussian kernel with standard deviation *σ*:

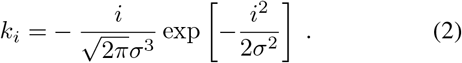

Eq. 1 corresponds to an approximate evaluation of the square magnitude of the intensity gradient, along the direction **n**. We choose *σ* = 2 pixels, a value which we determined empirically to ensure high enough smoothing of intensity profiles while maintaining a low computing cost. To obtain a force acting on a mesh node, one evaluates a finite difference:

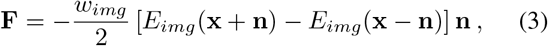

with *w*_*img*_ a factor modulating the weight of the contribution of the image energy relative to the intrinsic mesh forces. To calculate the energy at a point not located exactly at the center of a voxel, we use linear interpolation to evaluate the intensity *I*(**x**). This force is added to a force contribution intrinsic on the mesh, which depends on its curvature and the distance between nodes, to penalise surface bending and surface area [8] (Appendix B 1).

#### 2. Segmentation of cell membranes from original image data

To segment the membrane we use a “perpendicular intensity energy”. As the labelled membrane can be considered as a bright surface, we use an energy which attracts a mesh node to regions of high intensity. Considering a node at position **x** with unit normal vector **n**, defined as in section II A 1:

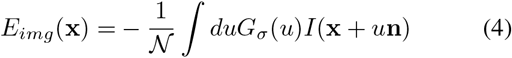

where *G*_*σ*_ is a one-dimensional Gaussian kernel with standard deviation of 𝒩 pixels, and is a normalisation factor. Eq. 4 corresponds to a convolution operation between the kernel *G*_*σ*_ and the intensity profile *I* evaluated along the normal **n**.

We then use the following force acting on a mesh node at position **x**, obtained by evaluating a discretized version of the gradient of the energy *E*_*img*_ in Eq. 4, along the normal to the mesh node **n**:

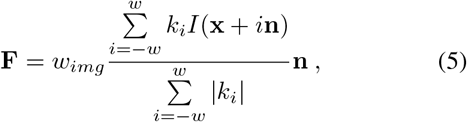

where *k*_*i*_ is defined in Eq. 2, and *w*_*img*_ is a factor modulating the weight of the contribution of the image energy relative to the intrinsic mesh forces. Here one can use a collection of manually created spheres to initialize a segmentation mesh, similar to what was done to segment cell nuclei; alternatively one can also use the nuclear mesh as initialisation and subsequently deform it to the membrane channel.

#### 3. Manual improvement of active mesh segmentation

A segmentation problem arises when the mesh does not stabilize to a steady state that suitably follows the contour of the object. Such a situation can be caused by image artefacts, poor initialization, or a poor choice of mesh deformation parameters. These issues can be addressed with DM3D by interactively editing meshes. Meshes can also be manually initialised more closely to the desired shape. Parameters *α, β* affecting the mesh evolution can be adjusted (see Appendix B 1 for a definition), and the resulting effect on mesh deformation can be observed directly within the plugin. Mesh deformation iterations can also be performed by modulating the weight *w*_*img*_ of the image energy relative to the intrinsic mesh energy. Reducing the role played by the intrinsic mesh energy allows the mesh to capture more prominent, irregular features of the cell nucleus or membrane. An additional tool is available within the DM3D plugin to manually edit meshes and deform them to the desired output.

### B. Neural network training

In order to train the neural network, we initially generated manual segmentations of cell nuclei and membranes for 3 time frames of Movie 2. Manual segmentation meshes are used to create training labels to train a 3D Unet [17]. As first described in Ref. [22], we modified the Unet architecture to predict 3 separate labels (Appendix C): i) a binary mask label that indicates all voxels contained within a mesh, ii) a binary label indicating the border of the binary mask, iii) a distance transform label with values ranging from 0 to 32. Labels are created for training by first generating a binary image (see Appendix B 3) from all of cell nuclei meshes or all of cell membrane meshes; in this binarisation, voxels which are contained within a mesh have value 1, and voxels outside have value 0. This binary image directly provides the mask label, while the binary label for the border are the edge voxels of the mask label. The distance transform is obtained by iteratively eroding the binary image in 3D, and labeling the eroded voxels with the current iteration depth value: the 0th depth eroded corresponds to border voxels, while voxels eroded at the next iteration have distance transform value of 1. We choose to saturate the distance transform value to 32, for ease of manipulation of images.

Two neural networks were trained using labels calculated (5) from the nuclei and membrane meshes respectively. Each network is trained to learn all three labels simultaneously by using a loss function that is the sum of three loss functions:

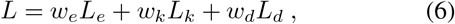

where *L*_*e*_, *L*_*k*_ and *L*_*d*_ are loss function for the border, mask and distance transform labels respectively, and *w*_*e*_, *w*_*k*_ and *w*_*d*_ are the corresponding weights in the total loss function. *L*_*e*_ and *L*_*k*_ are Sorensen-Dice coefficient loss functions, *L* = (|*TP* |+ 1)*/*(|*T*| + |*P*| + 1), and *L*_*d*_ is the Log-Mean-Square-Error *L*_*d*_ = log((*T −P*)^2^), with *T* the truth pixel values and *P* the network predicted pixel value. Neural network parameters can be adjusted to optimize the segmentation results. Here we found that setting the weights *w*_*e*_ = *w*_*k*_ = *w*_*d*_ = 1 in Eq. 6 led to acceptable results.

The distance transform contains in principle all of the information of the other two channels, so strictly speaking the membrane and mask channels do not need to be learned by the neural network. However, training the network to learn the membrane and mask labels helps to determine if the network is training properly. Incorrect learning of one of the training labels indeed likely indicates a problem with the training data.

### c. Obtaining nuclei segmentation meshes

To test the pipeline, we first used the network trained on nuclei labels to obtain nuclei segmentation meshes for all frames of Movie 2. To achieve this, we used the neural network to predict the distance transform of all frames of the movies. The predicted distance transforms are then turned into a binary image through a thresholding step, and continuous regions are labelled and filtered by size. We found that using a distance transform threshold of 1 did not allow to separate all nuclei, as some nuclei are close to each other. To address this, we selected a higher threshold value of 3, and use a region-growing or watershed algorithm to expand the detected regions, based on the distance transform image. The detected regions are then used to seed meshes, as follows: for each region, an approximately spherical mesh is generated by creating an isocahedral mesh, centered at the center of mass of the region, and subsequently subdividing the triangles of the mesh. Rays are cast from the center of mass of the region towards nodes of the spherical mesh. Each node is repositioned to the furthest voxel on the inner surface of the detected region that intercepts the corresponding ray [8]. The initialized mesh is then deformed by calculating the “perpendicular intensity energy” of the distance transform with a negative image weight (see section II A 1). This causes the mesh to be attracted to low values of the distance transform, away from the internal volume of the nucleus. A choice of positive and sufficiently large value of the parameter *α* (Appendix B 1) counteracts this effect by ensuring that the mesh tends to shrink and so wraps around the nucleus.

The step of mesh deformation is strongly affected by the quality of the neural network prediction. When the regions detected from the distance transform predicted by the neural network appear to correspond to a visible nucleus, the mesh deformation process reaches a steady-state. When a steady state cannot be found by the active mesh deformation algorithm, the mesh tends to shrink and can then be removed following detection of small volume meshes. This can indicate a false positive, where the neural network wrongly identifies a nucleus and the corresponding region needs to be removed. Failure of the mesh to converge to steady-state therefore acts as a filtering step.

To evaluate the segmentation results, we plotted the total number of cells over time. Fluctuations in cell count which do not correspond to cell division indicated that the network was failing to accurately segment some frames. For the first movie we segmented, Movie 2, a large number of mitosis events were causing the network to fail. We used DM3D to manually segment 5 additional frames (numbered 21-25) and trained the network using this additional data. After another iteration, we found that the later frames of the movie had some degradation in segmentation quality, due to a change in image quality. We therefore manually corrected a late time point (frame 132), and trained again the network including this frame. This step reduced the number of corrections required to segment late time points.

### D. Obtaining cell membrane segmentation meshes

To obtain cell membrane segmentation meshes, we use the predicted nuclei meshes to initialize active meshes, and deform them to the membrane distance transform predicted by the neural network trained using manually obtained membrane labels. We use a perpendicular intensity energy (Eq. 5), with a negative weight *w*_*img*_ to ensure that the mesh is converging to minima of the distance transform.

## III. RESULTS: SEGMENTING MAMMARY GLAND ORGANOIDS

### Test of fully automated pipeline on seen and unseen data

#### 1. Automated nuclei segmentation

To verify the quality of segmentation results, we compared fully automated segmentations to manually segmented validation data (Fig. 4). We used two sets of validation data: 9 “seen” 3D images which correspond to the training data taken from Movie 2, and 6 “unseen” 3D images which consist of single frames from Movie 1 to 6 (Fig. 2) that the network has not seen during training. The ground truth is a labelled image generated from manually segmented meshes, where each mesh is binarized and labelled with a unique number. A fully automated segmentation is generated as follows: the neural network is used to create a distance transform image for nuclei. Seed points are then determined from the distance transform based on a thresholding step with threshold value of 3. Seed points are used to initialise segmentation meshes for nuclei. These segmentation meshes are deformed using the “perpendicular intensity energy” of the distance transform, as described in section II C. Parameters for mesh iteration are given in Appendix B. The resulting meshes are used to create a fully automated labelled image, which can be compared to the ground truth labels.

**FIG. 4:**
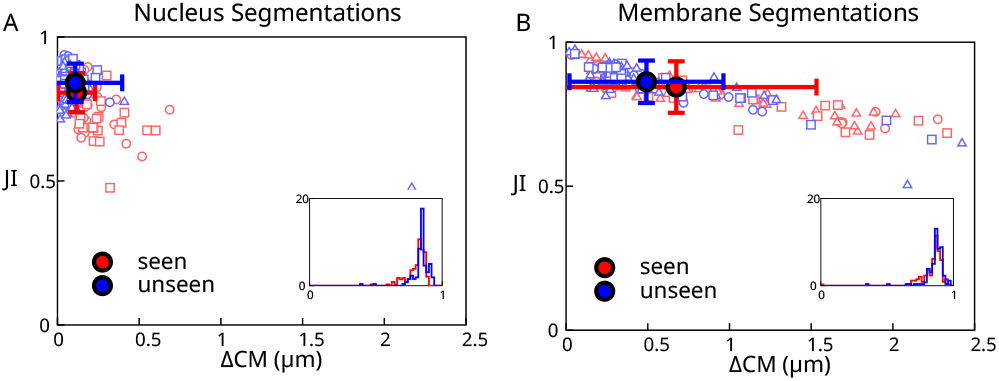
Analysis of automated segmentation quality. A,B) Scatter plot of best Jaccard Index (JI) versus the distance between the ground truth center of mass and the predicted center of mass (∆ CM) for cells from a “seen” and “unseen” dataset. A) Results of automated segmentation of cell nuclei at full resolution (voxels with side length 0.175 µm). A nucleus diameter is about 8 µm. B) Results of automated segmentation of cell membrane at full resolution. Insets: histogram of best Jaccard Index distributions. Individual data points outside of the plot range: A: 1/300, B: 2/300.

To measure the accuracy of the resulting automatic segmentation, we considered two metrics: the best Jaccard Index (JI), and the distance between the ground truth and predicted center of mass ∆CM (Fig. 4). The best Jaccard Index value for cell *i* JI_*i*_ is calculated for a given ground truth label *i* by calculating the Jaccard Index between *i* and each prediction label *j*, and finding the optimal value over prediction labels:

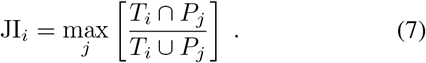

Here *T*_*i*_ denotes the set of voxels with the ground truth label *i, P*_*j*_ the set of voxels with predicted label *j, T*_*i*_ ∩ *P*_*j*_ is the size of the intersection between *T*_*i*_ and *P*_*j*_ and *T*_*i*_ ∪ *P*_*j*_ the size of the union, in number of voxels. The predicted cell that gives the maximum Jaccard Index is also used to calculate the distance between predicted and ground truth center of mass ∆CM_*i*_ for cell *i*.

In Fig. 4A we show a scatter plot in the space of values of (∆CM_*i*_, JI_*i*_) for each nucleus, as well as corresponding averages for all detected cells. This graph allows to visualise the accuracy of nuclei detection and reproduction of their shapes, using full resolution images to generate meshes for the nuclei. The pipeline achieves excellent results, with 98% of the unseen segmented cells with a JI above 0.7. Surprisingly, the pipeline achieves overall better results for unseen than from seen data. This may be because some of the “seen” dataset frames were selected because they caused segmentation issues due to cell mitosis or degraded image quality, while the “unseen” dataset was chosen arbitrarily and therefore has no comparable bias.

#### 2. Full resolution images, automated membrane segmentation

We then tested our pipeline on cell membrane segmentation. Here the automated membrane segmentation was obtained by adjusting meshes obtained from the automated segmentation of nuclei, using the predicted distance transform to the cell membrane, as described in section II D. Parameters for mesh iteration are given in Appendix B. The ground truth segmentation was obtained by manual edits of membrane segmentation meshes. Comparing the result of automated segmentation to the ground truth segmentation (Fig. 4B) shows that the automated segmentation is giving excellent results, although slightly less accurate than nuclei segmentations. This reflects additional difficulty in segmenting cell membranes: their shapes are generally more complex, for instance due to membrane appendages which are difficult to identify automatically at the imaging resolution achieved here.

### B. Full organoid segmentation over time

We then turned to full segmentation and tracking of the whole 24 hour organoid movies (Fig. 5 and 6). Using the automated segmentation steps described in section III A, we first obtained a fully automated segmentation of nuclei for all 6 movies.

**FIG. 5:**
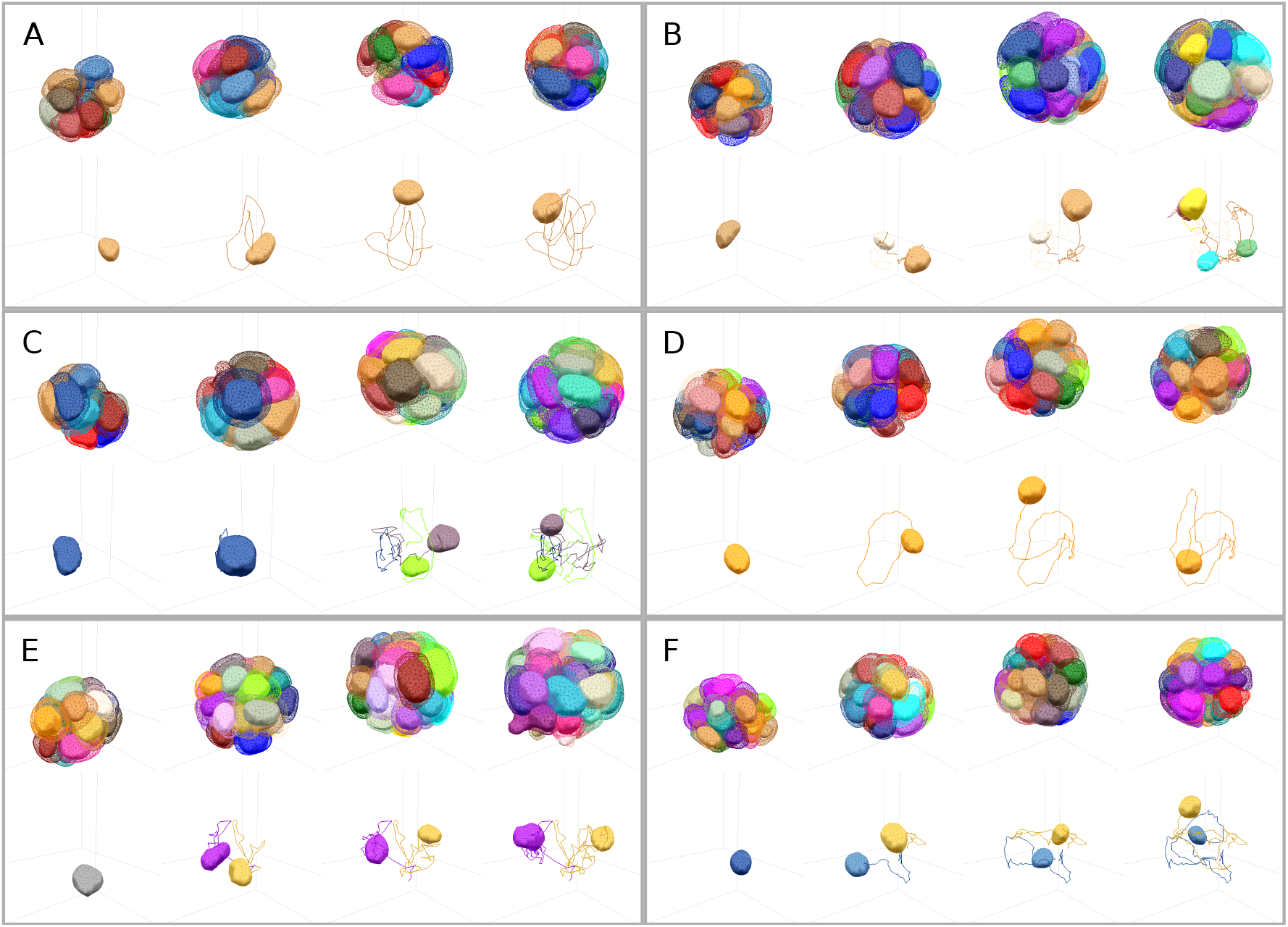
Segmentations results for cell membrane and cell nuclei for six different mammary gland organoids, segmented over 24 hours of growth. A-F) Within each box, segmentation meshes are shown at 0, 8, 16 and 24 hours for each organoid. Within each box, top row: solid volumes correspond to nuclei segmentation meshes and wireframes to cell membrane segmentation meshes. Bottom row: example trajectory of a cell nucleus and the nuclei of the cell progeny, during the movie.

**FIG. 6:**
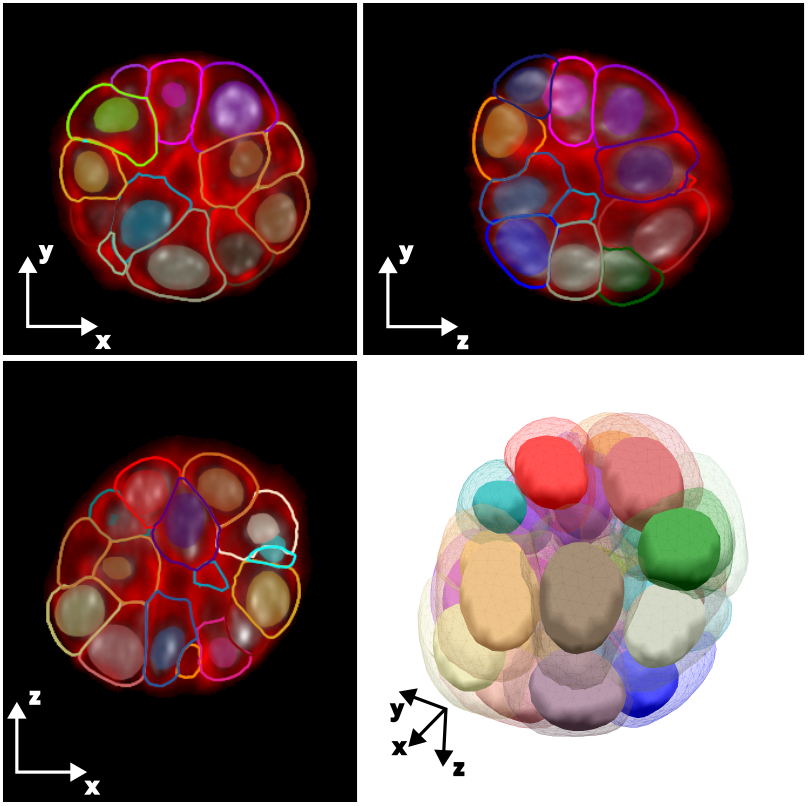
Cross-section and 3D view for one frame of one mammary gland organoid shown in Fig. 5. The cross-sections display overlay of nuclei (filled volumes) and membrane (wireframes) segmentation meshes on the original data (red: membrane dye, grey: DNA label).

To compare these results to a ground truth, we then manually corrected them. We proceeded as follows: segmentation of nuclei were used to track the cells over time by using a naive bounding box tracking algorithm (see Appendix B 6), and we quantified the cell count over time. Tracking errors and changes in cell count allow to find segmentation errors, when a nucleus appears or disappears, not due to cell division or death. Meshes were corrected by manually initialising a new mesh, deleting incorrect meshes, or splitting meshes that contain multiple nuclei. The corresponding dataset constitutes a new ground truth nuclei segmentation.

We then evaluated the detection accuracy of cell nuclei between this manually corrected dataset and the automated segmentation, for all time frames in the 6 organoid movies. To measure the detection accuracy, we mapped predicted to ground truth nuclei. We associate to each nucleus an axis-aligned bounding box, with axis aligned along the x,y,z directions of the image. We then compare the Jaccard indices of the bounding boxes of predicted and ground truth nuclei, as defined in Eq. 7. A predicted nucleus maps to the ground truth nucleus in the same frame with the highest JI value. We perform the symmetric operation and map ground truth nuclei to predicted nuclei. If a predicted nucleus and ground truth nucleus are singly mapped to each other, then we count the predicted nucleus as a True Positive (TP). When multiple predicted nuclei map to the same ground truth nucleus, then we count those predicted nuclei as False Positive (FP). If multiple ground truth nuclei map to a single predicted nucleus, or are not mapped at all, then these ground truth nuclei are counted as False Negative (FN). Better networks have a higher number of TP cells, and a smaller number of FP and FN cells. The corresponding results are reported in Table I. This showed that the automated procedure has an accuracy of *∼*90%, as evaluated by the fraction of TP cells.

**TABLE 1:** Detection accuracy for models at half and full resolution. Data corresponds to frames from all 6 organoids. *N* corresponds to the total number of segmented nuclei. The “half-resolution” model has been trained with 234 additional frames.

| Model | N | TP | FP | FN | TP/N |
| --- | --- | --- | --- | --- | --- |
| Full resolution | 18414 | 16630 | 182 | 1637 | 90.3% |
| Half resolution | 18414 | 18216 | 72 | 131 | 98.9% |

To visualise the outcome of the full organoid segmentation, we use corrected nuclei segmentation meshes to initialise membrane segmentation meshes. These meshes are then deformed according to a perpendicular intensity energy calculated with the neural network predicted distance transform to cell membranes. Here the procedure is fully automatic and no further correction is performed. The corresponding results for tracked nuclei and membrane meshes are plotted in Fig. 5. We used these nuclear segmentation results to evaluate cell motion in the organoids. All 6 organoids are highly dynamic, as quantified by histograms of cell velocity (Fig. 7A). Plotting the number of cells as a function of time also revealed signification variation in cell proliferation, with some organoids keeping a constant number of cells while others exhibit significant cell division (Fig. 7B).

**FIG. 7:**
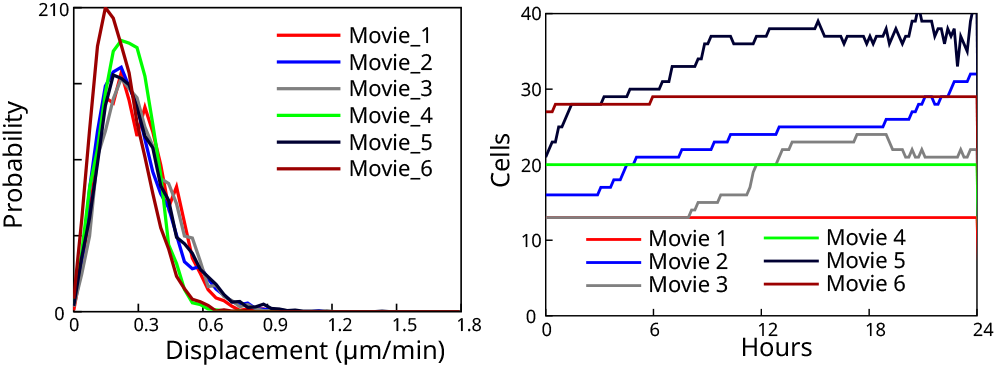
Quantifications associated to tracked nuclei for the 6 segmented organoids. A) Probability distribution of nucleus velocity, for each individual movie. B) Number of cells as a function of time.

### C. Reduction of image resolution and additional training

We then tested if the detection accuracy of cell nuclei could be improved by enlarging the training dataset. Incorporating a larger number of full resolution images in neural network training proved to be lengthy; therefore we resorted to half-resolution images. Training the network on half resolution images indeed requires 8 times less space, less memory requirement, and processing time.

To generate training data, we used nuclei segmented meshes from all 134 frames from Movie 2 and 100 frames from Movie 3 (excluding frames which are part of the “unseen” dataset described above), and trained a neural network on images at half resolution. We note that additional ground truth data in this larger dataset was manually curated with less accuracy than the original dataset used for initial training of the network. The network was trained over 116 epochs, during 10 days on a single Nvidia 3080 GPU workstation. For membrane segmentation data, we used the original training data consisting of 9 frames from Movie 2 at half resolution to train a neural network. Here the network was trained over 86 epochs, during 9 hours on a single Nvidia 3080 GPU workstation.

We then evaluated the quality of mesh segmentation resulting from this newly trained neural network. Comparing Fig. 8 to Fig. 4 shows that both the ∆CM prediction accuracy and the JI measurement are slightly worse with decreased image resolution, despite using an enlarged dataset. However, the prediction accuracy is still acceptable.

**FIG. 8:**
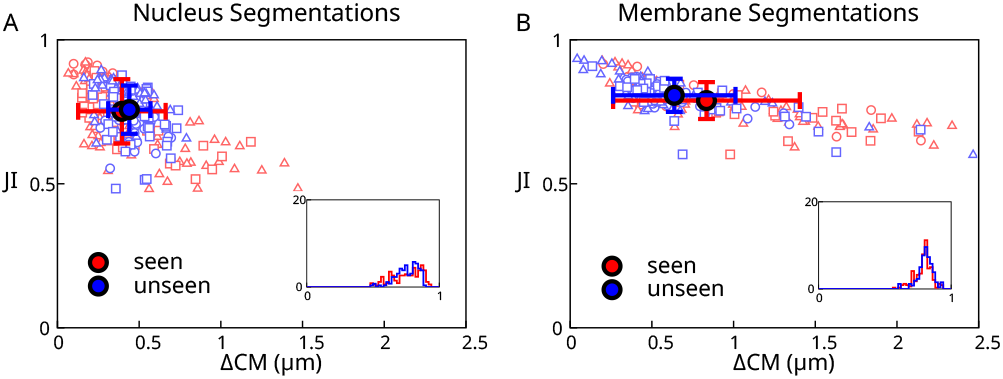
Analysis of automated segmentation quality at half-resolution, with a larger training dataset. A,B) Scatter plot of best Jaccard Index (JI) versus the distance between the ground truth center of mass and the predicted center of mass (∆CM) for cells from the same “seen” and “unseen” dataset as in Fig. 4 (here the training dataset is larger than the “seen” dataset). A) Results of automated segmentation of cell nuclei at half-resolution (0.350 µm voxels). B) Results of automated segmentation of cell membrane at half-resolution. Insets: histogram of best Jaccard Index distributions. Individual data points outside of the plot range: A: 0/300, B: 10/300.

We then evaluated the detection accuracy. Remarkably, training at half resolution with a larger dataset increased significantly the detection accuracy, reaching an excellent value of ∼ 99% (Table I). We think that this improvement can be attributed to the larger dataset used for training. We conclude that half-resolution images can be used for efficient and fast nuclei segmentation and tracking, while full resolution images can help with accurate nucleus and membrane segmentation. We note that the mesh representation is based on the actual size of the image volume, so that different scale images can be used with the same set of meshes.

### D. Comparison to StarDist

We then compared our segmentation results to outcomes obtained from the widely used Stardist software [23]. We generated StarDist labels using ground-truths labels from the “seen” set of images, as described in section III A 1. We trained two StarDist models, for the nucleus and membrane labels respectively, using the default parameters and with full resolution images. We use a provided default parameter of *N*_ray_ = 96 for the number of rays. We then tested the output of StarDist segmentation on the “seen” and “unseen” datasets (Fig. 9). We quantified the JI measurement and ∆CM prediction accuracy for nuclei and membrane, as was done using our pipeline (Figs. 4A,B and 9A,B). The comparison of these quantifications revealed that the StarDist segmentation outcome was slightly inferior to the result obtained with our pipeline, for both nucleus and membrane segmentation. However, we can not exclude that StarDist would not achieve better results by optimizing its parameters. For example, the number of rays determines the level of detail with which Stardist segment objects. We would expect that accurately segmenting cell membranes require more rays than segmenting nuclei. We note that in any case, a central advantage or our pipeline is the ability to easily manipulate and correct segmentation meshes, and use them to generate labels for further neural network training.

**FIG. 9:**
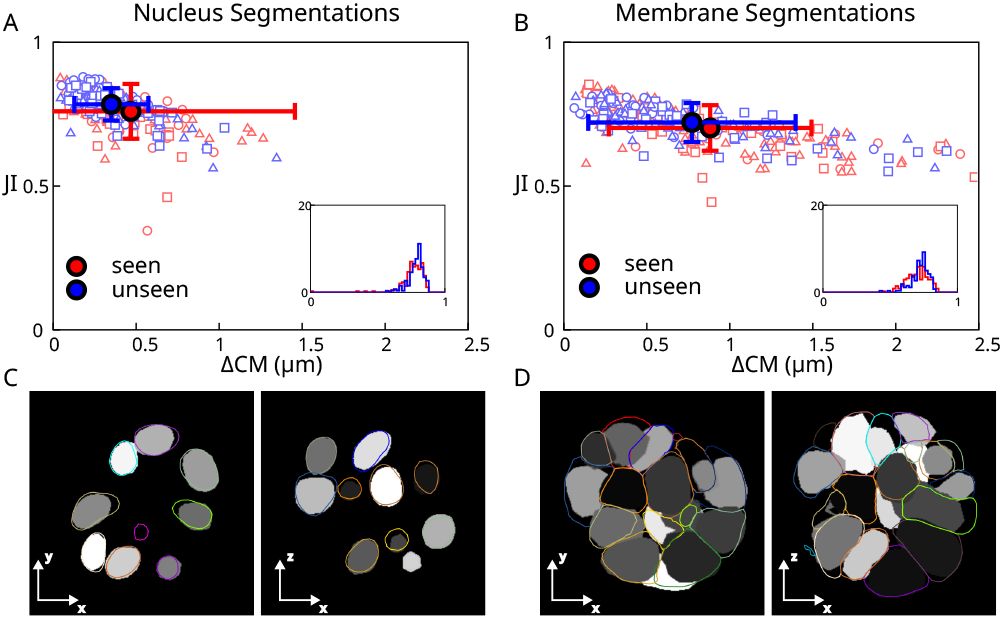
Analysis of segmentation quality with StarDist. A,B) Scatter plot of best Jaccard Index (JI) versus the distance between the ground truth center of mass and the predicted center of mass (∆CM) for cells from a “seen” and “unseen” dataset. A) Results of automated segmentation of cell nuclei. B) Results of automated segmentation of cell membrane. Insets: histogram of best Jaccard Index distributions. C) Representative example of nucleus prediction from StarDist, for two different planes of views. D) Representative example of membrane prediction from StarDist, for two different planes of views. In (C,D), grey regions correspond to StarDist predicted labels, coloured lines indicate ground truth segmentation meshes. Individual data points outside of the plot range: A: 2/300, B: 7/300.

## IV. RESULTS: SEGMENTING AGGREGATES OF MOUSE EMBRYONIC STEM CELLS

We then tested our methods on images from a different cell type, obtained with a different microscope. We applied our pipeline to a 10-frame movie of an aggregate of mouse embryonic stem cells (mESCs), imaged with a spinning disc microscope with 5 min time interval between frames (Fig. 10A). The resulting images have non-isotropic voxels, with a pixel size of 244 nm in the *x*-*y* plane and a 2 µm spacing between adjacent slices in the *z* direction. Because our neural network was initially trained on data with isotropic voxels, we interpolated the spinning disc images along the *z* axis to obtain modified images with isotropic voxels of size 244 nm. These modified images were then used for training the neural network and segmenting the images.

**FIG. 10:**
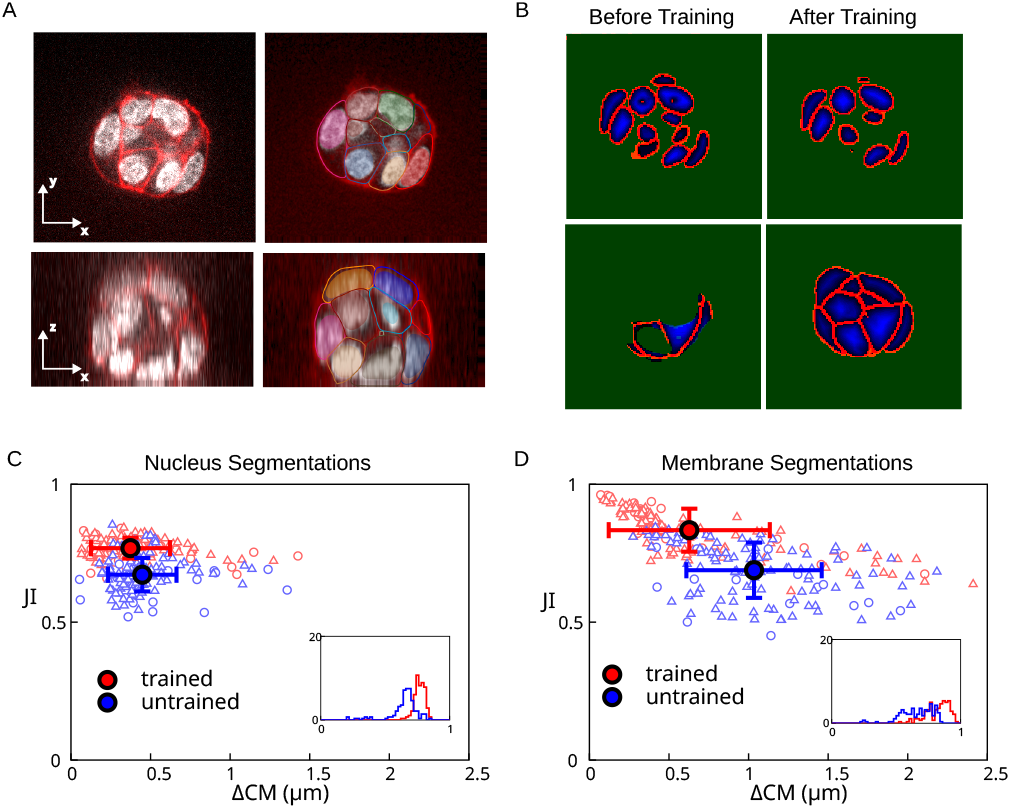
Mouse embryonic stem cell colony imaged on a spinning disc confocal microscope. A) Cross-sections of original image (left) with ground truth segmentation result overlaid (right). White: nuclear label, red: membrane label, other colors: contours of membrane segmentation meshes and filled regions of nuclei segmentation meshes. B) Cross-section of neural network output, before and after training the network on the spinning disc images. Top images: nuclei segmentation, bottom images: membrane segmentation. Colors correspond to different outputs of the neural network. Green: mask, red: border, blue: distance transform. Green mask label indicates background. C, D) Scatter plot of best Jaccard Index (JI) versus the distance between the ground truth center of mass and the predicted center of mass (∆CM), before (“untrained”) and after (“trained”) training of the network on 2 frames of a movie of the colony. C) Results of automated segmentation of cell nuclei. D) Results of automated segmentation of cell membrane. Insets: histogram of best Jaccard Index distributions. Individual data points outside of the plot range: C, trained: 0/140; C, untrained: 10/140; D, trained: 6/140, D, untrained: 18/140.

We first attempted to segment the mESCs aggregates with the network previously trained on mammary gland organoid aggregates. We found that the neural network provided outputs which were acceptable for nucleus segmentation, although some border voxels appeared inside the nuclei (Fig. 10B, “before training”). The neural network output for the membrane was however strongly underdetecting cells (Fig. 10B, “before training”). To improve on these results, we manually segmented two frames of the movie and retrained the neural network. We then generated fully automated segmentation meshes for nuclei and membranes for the 10 frames of the movie, as described in section III for mammary gland organoids. We manually corrected these meshes to obtain a ground truth segmentation. We then compared the results of the automated segmentation before and after training the network with two additional frames from the new dataset, against the ground truth (Fig. 10C, D). We found that retraining of the neural network significantly improved the segmentation accuracy, which reached values comparable to our results with mammary gland organoids, despite the limited size of the additional training data set (compare with Figs. 4 and 8). We conclude that we expect that our pipeline can be applied to datasets coming from different microscopes and different cell types.

We observed that mESCs aggregates often have closely spaced nuclei, making their segmentation challenging. We found that the ability of the neural network to predict the distance transform aids in separating nuclei from one another, as the predicted distance transform can be used to identify the center of the nuclei.

## V. DISCUSSION

We showed that using a neural network is an effective way to initialise and deform active meshes on a large number of images. We found that combining active meshes segmentation with a deep learning neural network has several advantages. Notably, active meshes provide a direct and intuitive understanding of the origin of successful or failed segmentation, in contrast to neural network predictions. Relaxation of a mesh to a steady-state generally indicates that the image is of high enough quality for segmentation to succeed. If the mesh does not reach a steady-state, manual inspection of the image helps the user to understand the origin of failure. For instance, in the organoids we have segmented, we have found that automated nucleus segmentation by the neural network could fail because of nuclear dye accumulation artefacts, which could attract the nucleus segmentation mesh, or because of nuclear envelope breakdown during mitosis, as a well-defined nucleus is not visible. Manual mesh initialisation and subsequent mesh deformation allows to correct for these issues. In addition, retraining the neural network after mesh correction allows to obtain a predicted distance transform which improves on these issues. Overall, the combination of neural network prediction with active meshes allows for efficient manual curation and post-processing of the segmentation data and improvement of neural network prediction. Following manual curation of segmentation data, retraining of the network improves the outcome of automated segmentation. As Table I indicates, we could improve the accuracy in nuclei detection from *∼*90% to *∼*99% by manually correcting 234 frames and retraining the network; showing the importance of using a neural network in our pipeline. Possibly, repeating these steps of manual correction and network training may allow to further increase this accuracy.

When considering a new dataset to segment, manually segmenting with active meshes also allows to directly test whether the image quality is sufficient for segmentation. This step can be more revealing than directly segmenting a new dataset with a neural network, where segmentation failure could arise from from inadequate parameters within the neural network, but also from insufficient image quality.

In addition, mesh segmentations are independent of image resolution; this can be useful for locally downloading lower resolution images, or for generating training data at different resolutions.

We also note that the algorithm used to deform the active mesh can be applied to the image directly, instead of the distance transform prediction returned by the neural network. This can in principle ensure that the final segmentation result is independent of the parameters of the neural network and its training history.

Using a neural network also alleviates known drawbacks to active meshes: that an initialisation seed has to be found by hand, and that deformation parameters need to be adjusted for different image conditions. Indeed, in addition to providing with a high quality initialisation of the active mesh, the neural network effectively removes noise and, through the prediction of a distance transform, adjusts signal levels, such that a good set of active mesh parameters will work over a larger range of data qualities.

In this study we have considered two datasets where cells have relatively regular shapes. Our pipeline might have to be adapted to segment more complex cell shapes, possibly by adjusting deformation and re-meshing parameters of active meshes.

The DM3D interactive plugin used in this study was built around an active mesh deformation method [8], introduces handling of multiple meshes in the same time frame, steric interaction between meshes, and a remeshing algorithm (Appendix B), so that the plugin is adapted to organoid segmentation. The plugin can also share formats, and produce segmentations from a variety of image sources. The plugin also works with virtual image stacks and can be used with or without a 3D display; this makes it practical to work both locally or on remote computers. It is also effective for monitoring segmentations at different points in the pipeline. In an effort to make our plugin more accessible we have added ways to export meshes as other 3D mesh formats, as TrackMate [24] files to apply more advanced tracking algorithms, or as integer labelled images. In addition to describing the DM3D plugin, we have reported the development of a new 3D-Unet based segmentation approach that works in conjunction with the DM3D tool. Neural networks and the DM3D plugins are available as described in Appendix A. We provide a tutorial which can be used to analyse example data with 6 frames and a few cells. Generation of neural network prediction and active mesh evolution take a few minutes on a standard laptop to generate for images used in this tutorial.

We believe that the combination of active meshes and neural network offers a flexible and efficient way of segmenting 3D image data, and we hope that our tool will prove valuable for the scientific community.

## Supporting information

Movie 6 segmentations aligned by group rotations

Movie 6 segmentations aligned by center of mass

## Acknowledgements

MBS and GS acknowledge support from the Francis Crick Institute, which receives its core funding from Cancer Research UK (FC001317), the UK Medical Research Council (FC001317), and the Wellcome Trust (FC001317). MBS, HS, GS, CD, AB acknowledge support from a grant to GS, CD, AB from the Engineering and Physical Sciences Research Council (EP/T003103/1). AC acknowledges funding from the Wellcome Trust (201334/Z/16/Z) and Utrecht University. G.S. thanks Kai Dierkes for discussions on neural networks.

## Author contributions

MBS, HS and JA performed research with supervision from AB, CD, and GS. MBS developed and applied the segmentation pipeline, JA prepared mammary gland organoids and HS performed imaging. AC acquired the data in Fig. 9. MBS and GS wrote the manuscript with inputs from CD.

## Appendix A

**Code and data availability**

This project is composed of two open source projects available on github: DM3D an interactive plugin for creating and deforming meshes, and ActiveUnetSegmentation a tensorflow implementation of a 3D Unet available at https://github.com/PaluchLabUCL/DeformingMesh3D-plugin and https://github.com/FrancisCrickInstitute/ActiveUnetSegmentationrespectively. DM3D is also distributed as a Fiji plugin by using the Fiji update site, https://sites.imagej.net/Odinsbane. Additional documentation and usage examples can be found at https://franciscrickinstitute.github.io/dm3d-pages/. A detailed tutorial for the DM3D plugin, with example data, can be found at https://franciscrickinstitute.github.io/dm3d-pages/tutorial.html. Additional data and trained neural networks used in this study can be found at https://zenodo.org/record/7544194.

## Appendix B

**Details of the DM3D plugin**

In this Appendix we provide details of the active mesh DM3D plugin.

### 1. Mesh iteration

As described in Ref. [8], a mesh node *i* with position 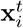 at pseudotime *t* of mesh evolution, evolves according to the following equation:

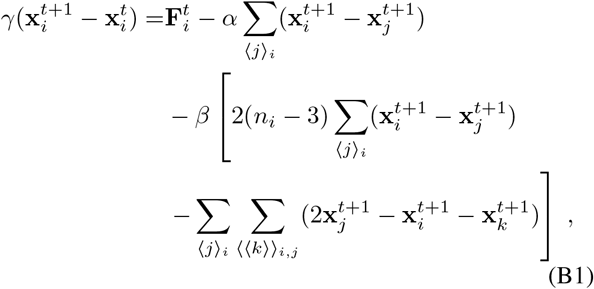

where ⟨ *j* ⟩ _*i*_ denotes the set of nodes *j* directly connected by an edge to node *i, n*_*i*_ is the number of nearest neighbours of node *i*, and ⟨⟨*k* ⟩ _*i,j*_ denotes the set of nodes *k* neighbours of *j* which are not neighbours of *i*. 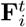 is an additional force which is obtained from Eq. 3 or 5. *α* and *β* are mesh evolution parameters which can be adjusted.

For automated segmentation of nuclei based on the predicted distance transform, we use the “perpendicular intensity energy” with *α* = 1, *β* = 0.1, *γ* = 1000 and *w*_*img*_ =*−* 0.05, and perform 100 iterations of each mesh. The same parameters were used for automated segmentation of membrane based on the distance transform, except with *w*_*img*_ = − 0.1, 800 iterations of each mesh, and intermediate steps of automatic remeshing with minimum length 0.75 µm and maximal length 1.6 µm.

### 2. Remeshing

We use a remeshing algorithm which splits long edges (larger than a threshold) and remove short edges (smaller than a threshold). This allows meshes to deform in an unconstrained manner. Remeshing is performed by first sorting each edge by length. The longest edge is then split in two and the two adjacent triangles are replaced by four new triangles. The process is iterated going through edges by decreasing order of length.

Once all of the long edges have been split through this process, short edges are removed if a number of conditions are satisfied. One denotes *i* and *j* the nodes connected by the edge. One then finds the sets of neighbouring nodes that share an edge with nodes *i* and *j*. If the neighbours of *i* and the neighbours of *j* have exactly 2 nodes in common, denoted *k* and *l*, then the connection is removed if both *k* and *l* have strictly more than 3 edges connected to them. If the two sets of neighbours have 3 nodes or more in common, the edge is not removed. These criterions allow to prevent meshes with problematic topologies. After removal of the edge, a new node replacing nodes *i* and *j* is generated at the mid point between the nodes *i* and *j*; edges previously connected to *i* and *j* are connected to the new node, and duplicate edges are removed.

The remeshing algorithm significantly improves the quality of meshes, by allowing them to deform to more exotic shapes with better distributions of triangles.

### 3. Binarizing a mesh

We use the following procedure to obtain a binary image from segmentation meshes, with label 0 indicating voxels outside of segmentation meshes and label 1 indicating voxels inside meshes. For each *y, z* value in the 3D image we cast a virtual ray through the mesh, going along the *x* axis. As one progresses along the *x* axis of the image, a topological depth is iterated, starting from value 0. When the ray crosses the mesh, the topographical depth increases by 1 if the scalar product between the normal and the unit vector giving the direction of the ray is negative. Indeed, meshes are defined with normal vector of triangles pointing towards the outside. Conversely the depth decreases by 1 if the scalar product is positive. The voxels are then scanned across and are determined to be inside of the mesh if they coincide with a region of positive depth, and outside if they coincide with a region of zero or negative depth.

### 4. Modified distance transform

The modified distance transform is found by iteratively eroding the binary representation of a segmentation mesh. A distance transform image is initialized with voxels with value At first, all positive binary voxels that are neighbouring a 0 valued voxel in the binary image are allocated a distance transform value of 0. A new eroded binary image is obtained by setting these voxels to 0. All positive binary voxels that are neighbouring a 0 value voxel are then allocated a distance transform value of 1, and a new eroded binary image is obtained by setting these voxels to 0. This process is iterated up to a distance transform value of 32; remaining positive binary voxels are allocated a distance transform value of 32.

### 5. Steric energy

To help with semi-automated mesh-based segmentation, we have introduced a steric energy between active meshes. Several meshes can be evolved simultaneously according to Eq. B1, with an additional contribution to the force 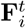 that minimizes a steric interaction energy. This method can be used to help deform segmentation meshes, when a feature of the image prevents them from deforming properly if unconstrained. The extra contribution to the force 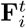is calculated based on the penetration depth of mesh points into neighbouring meshes. Here we have not used this tool for automated segmentations.

### 6. Tracking algorithm

Tracking is performed by bounding box Jaccard index detection. The axis-aligned bounding box of each mesh is calculated, and the bounding boxes of successive frames are used to calculate the Jaccard index. Cells with the highest Jaccard index between successive frames are mapped to each other. After this first pass tracking algorithm is used, manual tracking error correction can be performed. Tracking errors can be found notably from large displacements, or tracks ending abruptly.

## Appendix C

**Unet modification**

We trained a Unet network with the architecture described in Ref. [17], with the following modifications: we use 3 separate convolution output layers instead of a single one. Two ouput layers for the mask and distance transform are obtained from the network at depth 0, while the output layer predicting the object border is obtained from the network at depth 1. We use different activation functions for the 3 output layers: sigmoid activation for the mask and border labels, and ReLu activation for the distance transform.

## Appendix D

**Material and methods**

### 1. Mouse Models

Mice were bred and maintained at the Biological Research Facility of the Francis Crick Institute and the Biological Services Unit of the Institute of Cancer Research. MMTV-PyMT, mTmG and LifeAct-GFP mice were described before [25– 27]. Mice were kept in individually ventilated cages at 21^*◦*^C and fed ad libitum. Mouse husbandry and euthanasia for tissue collection was performed conforming to the UK Home Office regulation under the Animals (Scientific Procedures) Act 1986 including the Amendment Regulations 2012. Ear biopsies were sampled for genotyping. All mice were culled by cervical dislocation and confirmation of cessation of circulation.

### 2. Organoid culture

Organoids were established from healthy mammary glands or mammary tumours of females of 10-12 weeks of age. Mice were humanely culled by cervical dislocation and tissue dissected in aseptic conditions. Mammary glands were digested in 30 µg/mL collagenase (SIGMA-Aldrich) in DMEM/F12 (GIBCO) using a MACS dissociator (Miltenyi) at 37^*◦*^C for 20 minutes. Digested tissue was transferred to a 15 mL centrifuge tube and centrifuged at 3,000 rpm, supernatant discarded, and pellet resuspended in 1 mL red blood cell lysis buffer (SIGMA-Aldrich) for 3 minutes. 9 mL of DMEM/F12 were used to stop the red blood cell lysis and tubes were centrifuged in the same conditions. Supernatant was discarded and pellet resuspended in 1 mL of 1x trypsin (GIBCO) and incubated at 37^*◦*^C for 3 minutes. 5 mL of DMEM/F12 with 10% FBS (GIBCO) were used to stop the trypsinization reaction and tubes were centrifuged. Supernatant was discarded and pellet resuspended in 5 mL DMEM/F12, passed through a 70 µm filter and centrifuged again. Supernatant was discarded and pellet resuspended in Matrigel™ (Corning) and plated in 25 µL domes, one dome per well of a 24-well plate (Costar) for maintenance. Domes were polymerized at 37^*◦*^C for 30 minutes and covered in mouse mammary gland organoid media consisting of 50 ng/mL EGF (Preprotech), 100 ng/mL FGF (Preprotech), 4 µg/mL heparin (SIGMA-Aldrich), 1X B27 (GIBCO), 1X N2 (GIBCO), 1X penicillin/streptomycin (GIBCO), 10 mM HEPES (GIBCO) and L-glutamine-containing DMEM/F12 (GIBCO). Organoids were maintained at 5% CO2 and 20% O2 at 37^*◦*^C. For maintenance, weekly organoid splitting was performed by washing the Matrigel™ dome in 500 µL PBS, digestion in 300 µL TrypLE (GIBCO) for 10 minutes, dilution of TrypLE in 700 µL DMEM/F12, centrifugation at 3,000 rpm for 3 minutes, discarding supernatant, and resuspending pellet of cells in Matrigel™. For microscopy, cells were disaggregated in TrypLE as described before, counted using Trypan Blue and an automatic cell counter, resuspended in Matrigel™ and plated in a 96-well plate (Cellvis) in a 30 µL disc per well. Matrigel™ was polymerized for 30 minutes at 37^*◦*^C and wells topped up with 200 µL organoid media containing 3*µ*M SiR-DNA dye (Spirochrome). For position registration, 1:20 TetraSpeck™ 0.1 µm beads (ThermoFisher) were resuspended in Matrigel™, plated in 30 µL discs, and topped up with 200 µL organoid media.

### 3. Dual-view oblique plane microscopy (dOPM)

For time-lapse imaging of multiple organoids in parallel in a multi-well plate format, a dual-view oblique plane microscope (dOPM) was used, which was a modified version of the system reported in reference [20]. Briefly, the system is a type of light-sheet fluorescence microscope (LSFM) [28] that employs a single-objective for sample illumination and fluorescence detection [29] designed for multi-view single plane illumination microscopy (mSPIM) [30]. This type of microscope is suitable for fast 3D imaging with low light dose and reduced sample-induced image artefacts, so can be applied to timelapse imaging of multiple live organoids in parallel. For this work, the dOPM configuration reported in [20] was modified to operate with a Nikon 1.2 NA 60x water immersion objective as the primary microscope objective, which has a higher numerical aperture than the original design based around a Nikon 1.15 NA 40x water immersion objective [20]. The dOPM system was configured to record 3D image datasets in sample space from the perspectives of overlapping views that are rotated by ±35 degrees relative to one another about the optical axis of the primary microscope objective. Organoids were imaged every 11 minutes for 24 hours totalling 135 time points. From the two dOPM view’s perspectives, the acquired 3D image data per time point consisted of optically sectioned images spaced 0.6 µm apart covering a scan range of 90 µm. Each image plane was 450 pixels in width and height and the pixel size in sample space was 0.175 µm in each dimension to cover a field of view of 140 µm in each dimension. The illumination light-sheet used had a calculated full width at half maximum of 3 µm at the waist in the sample plane.

To image fluorescence from tdTomato labelled membrane, a 561 nm laser was used for fluorescence excitation and a 600/52 nm (central wavelength/band pass) emission bandpass emission was used for detection. To image fluorescence from nuclear SiR-DNA, a 642 nm laser was used for fluorescence excitation and a 698/70 nm (central wavelength/band pass) emission bandpass emission was used for detection.

For each spectral channel the information from the 3D dataset of each dOPM view was combined into a single 3D dataset by using a fusion routine in the Multiview fusion plugin available in ImageJ [31]. This routine required registration information to correctly co-register the two dOPM view. This information was determined from dOPM datasets of 3D samples of beads suspended in Matrigel which was included in the multiwell plate assay as discussed in section D 2 – see [31] for details of the bead-based co-registration method. Following fusing, the 3D datasets were converted to tiff stacks for segmentation.

#### 4. Culture and imaging of mouse embryonic stem cells

Mouse embryonic stem cells (E14 cells stably expressing H2B-RFP [32] were cultured as described in [33] on 0.1% gelatin in PBS (in N2B27+2i-LIF + penicillin and streptomycin, at a controlled density 1.5 - 3.0 *×* 10^4^ cells cm^*−*2^) on Falcon flasks and passaged every other day using Accutase (Sigma-Aldrich, #A6964). They were kept in 37^*◦*^C incubators with 7% CO2. Cells were regularly tested for mycoplasma.

The culture medium was made in house, using DMEM/F-12, 1:1 mixture (Sigma-Aldrich, #D6421-6), Neurobasal medium (Life technologies #21103-049), 2.2 mM L-Glutamin, home-made N2 (see below), B27 (Life technologies #12587010), 3 µM Chiron (Cambridge Bioscience #CAY13122), 1 µM PD 0325901 (Sigma-Aldrich #PZ0162), LIF (Merck Millipore # ESG1107), 0.1 mM *β*-Mercaptoethanol, 12.5 µg mL^*−*1^ Insulin zinc (Sigma-Aldrich #I9278). The 200 X home-made N2 was made using 8.791 mg mL^*−*1^ Apotransferrin (Sigma-Aldrich #T1147), 1.688 mg mL^*−*1^ Putrescine (Sigma-Aldrich #P5780), 3 µM Sodium Selenite (Sigma-Aldrich #S5261), 2.08 µg mL^*−*1^ Progesterone (Sigma-Aldrich #P8783), 8.8% BSA.

For colony imaging, the cells were plated on 35 mm Ibidi dishes (IBI Scientific, #81156) coated with gelatin the day before the experiment, and imaged on a Perkin Elmer Ultraview Vox spinning disc (Nikon Ti attached to a Yokogawa CSU-X1 spinning disc scan head) using a C9100-13 Hamamatsu EMCCD Camera. Samples were imaged using a 60X water objective (CFI Plan Apochromat with Zeiss Immersol W oil, Numerical Aperture 1.2). The samples were imaged acquiring a Z-stack with ∆Z = 2 µm every 5 minutes.

